# From Circles to Signals: Representation Learning on Ultra-Long Extrachromosomal Circular DNA

**DOI:** 10.1101/2025.11.22.689941

**Authors:** Jien Li, Zhenke Liu, Ziqi Zhang, Jiaqi Zhang, Ritambhara Singh

**Affiliations:** Department of Molecular Biology, Cell Biology and Biochemistry, Brown University, Providence, RI 02906; Department of Computer Science, Brown University, Providence, RI 02912; Center for Computational Molecular Biology, Brown University, Providence, RI 02912

## Abstract

Extrachromosomal circular DNA (eccDNA) is a covalently closed circular DNA molecule that plays an important role in cancer biology. Genomic foundation models have recently emerged as a powerful direction for DNA sequence modeling, enabling the direct prediction of biologically relevant properties from DNA sequences. Although recent genomic foundation models have shown strong performance on general DNA sequence modeling, their application to eccDNA remains limited: existing approaches either rely on computationally expensive attention mechanisms or truncate ultra-long sequences into kilobase fragments, thereby disrupting long-range continuity and ignoring the molecule’s circular topology. To overcome these problems, we introduce eccDNAMamba, a bidirectional state space model (SSM) built upon the Mamba-2 framework, which scales linearly with input sequence length and enables scalable modeling of ultra-long eccDNA sequences. eccDNAMamba further incorporates a circular augmentation strategy to preserve the intrinsic circular topology of eccDNA. Comprehensive evaluations against state-of-the-art genomic foundation models demonstrate that eccDNAMamba achieves superior performance on ultra-long sequences across multiple task settings, such as cancer versus healthy eccDNA discrimination and eccDNA copy-number level prediction. Moreover, the Integrated Gradient (IG) based model explanation indicates that eccDNAMamba focuses on biologically meaningful regulatory elements and can uncover key sequence patterns in cancer-derived eccDNAs. Overall, these results demonstrate that eccDNAMamba effectively models ultra-long eccDNA sequences by leveraging their unique circular topology and regulatory architecture, bridging a critical gap in sequence analysis. Our codes and datasets are available at https://github.com/zzq1zh/eccDNAMamba.

## 1 Introduction

Extrachromosomal circular DNA (eccDNA) is widespread in eukaryotic cells and often derives from the genome [1]. Specifically, eccDNAs possess a covalently closed, head-to-tail circular structure, introducing *wrap-around dependencies* that play important roles in DNA rearrangement. EccDNA is especially prominent in cancer, with sequence lengths that often exceed 500 kilobases and can even reach megabase scale [2]. These ultra-long molecules frequently harbor oncogenes and distal regulatory sequences that influence tumor evolution, therapeutic resistance, and intratumor heterogeneity [3, 4]. Recently, foundation models trained directly on large-scale DNA sequences have emerged as powerful tools for learning the underlying functional structures of the genome. Despite their success in DNA sequence modeling, existing genomic foundation models still face significant challenges when modeling the ultra-long and circular structure of eccDNA molecules. First, in cancer cells, the eccDNA sequences often span tens of kilobases to several megabases [5] [6], containing entire genes and regulatory regions. Modeling such ultra-long sequences requires a framework that can efficiently capture long-range contexts across extensive genomic sequences. However, the state-of-the-art attention-based foundation models, such as DNABERT-2 [7] and Nucleotide Transformer [8], while powerful for short sequences, scale poorly with input length due to the quadratic complexity of the attention mechanism. This computational bottleneck makes their direct application to ultra-long eccDNA infeasible. In practice, these models typically rely on truncating input sequences into kilobase fragments, disrupting the continuity of the original molecule and eliminating critical long-range contextual information, potentially leading to incomplete or misleading downstream analyses.

To meet the long-range context modeling requirement, other architectures replace attention with more computationally efficient operators. But these approaches still break the circular continuity at the head–tail junction of eccDNA. For example, HyenaDNA and its variant [9, 10] combine information across far-apart sequence positions without computing every intermediate interaction to achieve efficient scaling on very long sequences, yet its unidirectional computation limits the ability to model circular topology and loses dependencies spanning the head-to-tail connection. Caduceus [11] adopts bidirectional state-space models (SSMs) that read sequences step-by-step while maintaining a running summary to capture long-range patterns efficiently with base-level (per-nucleotide) tokenization, yielding fine-grained, symmetric context at single-base resolution. While SSMs provide linear-time sequence processing, making them well-suited for long inputs, the per-base tokenization expands the sequence length drastically. As a result, even with linear computation complexity, memory constraints often necessitate sequence truncation. Consequently, no existing method simultaneously achieves efficient modeling of ultra-long eccDNA sequences while preserving their intrinsic circular topology.

In this paper, we aim to resolve these modeling gaps by introducing eccDNAMamba [12], the first bidirectional state-space models tailored to eccDNA. eccDNAMamba first applies Byte-Pair Encoding (BPE) [13, 14] to compress repetitive nucleotide patterns into compact tokens. It then fuses forward and reverse Mamba-2 [15] passes to obtain full-context representations in linear time. Finally, through a circular augmentation implemented by appending a short snippet of the sequence’s beginning to its end, eccDNAMamba explicitly preserves the circular junction and wrap-around dependencies. This augmentation ensures topology-awareness while enabling eccDNAMamba to learn the representation of eccDNA sequences spanning megabases without truncation efficiently. We also systematically define and evaluate the core predictive tasks that make eccDNA modeling both biologically meaningful and computationally challenging.

To evaluate the effectiveness of our model, we preprocess data from CircleBase, eccDNAdb, and eccDNA Atlas [5, 6, 16] and establish the EccDNA Multi-Task Benchmark. This benchmark spans two biologically relevant classification tasks. The cancer-versus-healthy eccDNA discrimination task assesses whether an eccDNA sequence originates from cancer tissues. The copy-number level prediction task evaluates the degree of circular DNA amplification [2, 17]. Our results show that eccDNAMamba outperforms existing state-of-the-art foundation models for all these tasks. For cancer-versus-healthy eccDNA discrimination, eccDNAMamba surpasses the best-performing baseline. For eccDNA copy-number level prediction, eccDNAMamba exceeds the strongest baseline by 6.4% on average. Beyond accuracy, eccDNAMamba offers linear-time scaling and a low, stable memory footprint, enabling efficient eccDNA modeling.

We further demonstrate that eccDNAMamba can provide biological insights on ultra-long eccDNA sequences, unlike existing eccDNA tools that either overlook sequence-level modeling [18] or are restricted to short inputs [19]. Using Integrated Gradients (IG), our eccDNAMamba model highlights which regions of the ultra-long eccDNA sequences the model focuses on, therefore enabling direct biological insight into how model predictions relate to circular topology and regulatory features. These analyses reveal that eccDNAMamba focuses on regulatory and transposable elements that amplify oncogenic programs. Motif discovery on high-attribution regions recovers known oncogenic transcription factor families and uncovers novel sequence patterns of cancer eccDNA. The model effectively leverages the unique topology and regulatory architecture of eccDNA, providing a representation-learning framework for predicting and analyzing properties of eccDNA from sequence for the biology community.

## 2 Method

### Overview

We frame eccDNA analysis as a sequence modeling and prediction task: given an ultra-long eccDNA sequence, the goal is to learn representations that capture both its internal architecture and its dependencies, and to use these representations to support downstream predictions. (Fig. 1). Each raw DNA sequence, composed of the four nucleotide bases (A, T, C, and G), is first tokenized using Byte-Pair Encoding (BPE), which merges frequent nucleotide patterns into compact tokens to reduce sequence length while preserving biological meaning (Sec. 2.1). The model then applies circular augmentation by appending the first 64 tokens of each sequence to its end (Sec. 2.2), explicitly exposing the head–tail junction and enabling the model to learn wrap-around dependencies (Sec. 2.3). The model employs Mamba-2 encoders to process the augmented sequence from both directions: one pass moving forward and the other moving backward. The information gathered from the two directions is then integrated into a single, unified representation of the entire eccDNA sequence (Sec. 2.3).

**Figure 1:**
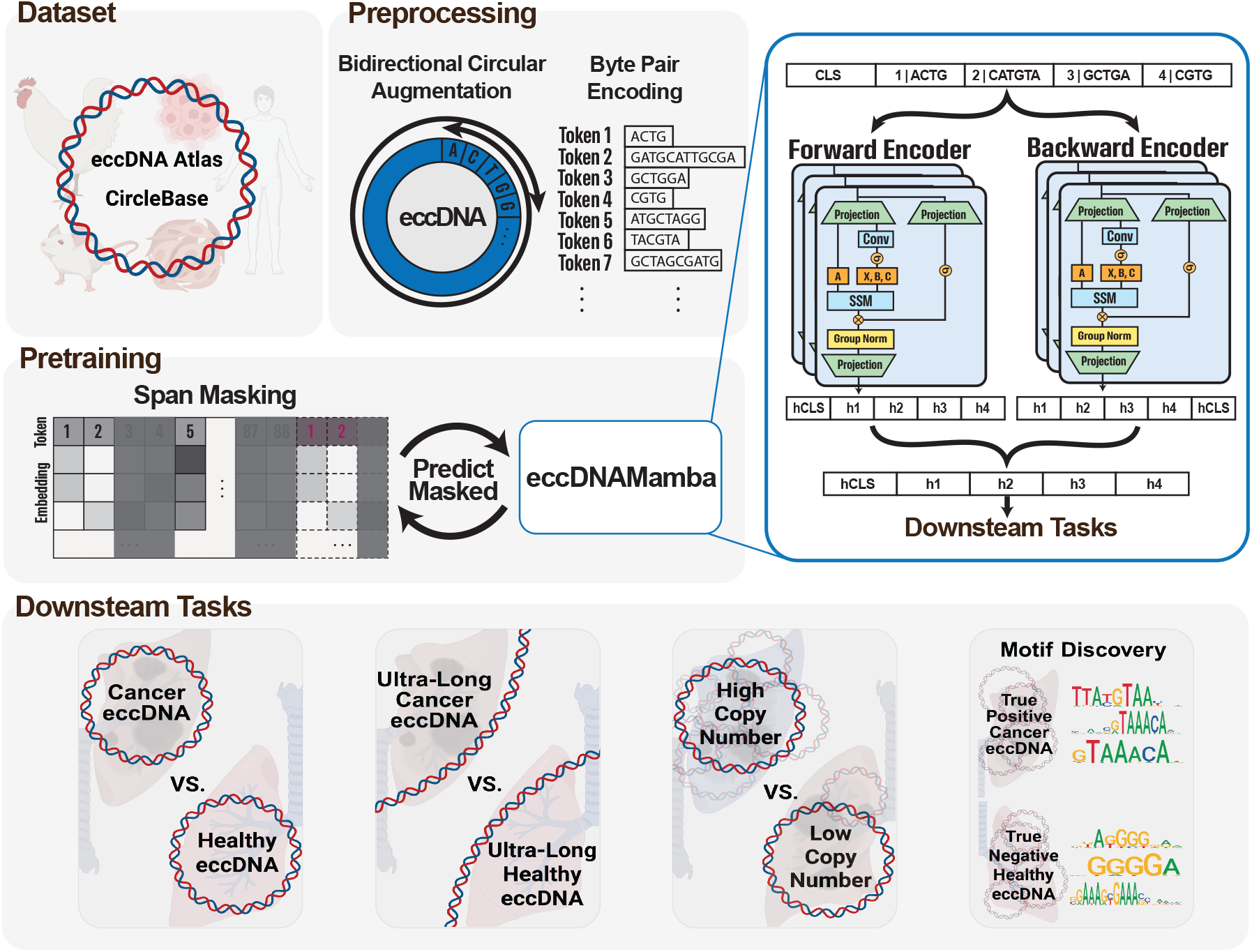
eccDNAMamba processes eccDNA through a topology-aware pipeline combining BPE tokenization, circular augmentation, and bidirectional Mamba-2 encoding to capture wraparound dependencies. The forward and reverse representations are fused via a shared MLP to generate unified sequence embeddings. During pretraining, a span-masked language modeling objective enables the model to learn long-range and circular dependencies for downstream genomic tasks.

During pretraining, eccDNAMamba is optimized with a span-masked language modeling objective, where random contiguous spans of tokens are masked, and the model learns to reconstruct the missing segments from the surrounding context (Sec. 2.4). This pretraining scheme encourages the model to capture long-range dependencies and circular topology in an unsupervised manner, providing general-purpose representations that can later be fine-tuned for downstream tasks such as cancer eccDNA classification and copy-number level prediction. We describe all these steps below.

### 2.1 Efficient Tokenization

To process genomic sequences with modern deep learning models, we need to convert raw nucleotide sequences into discrete tokens, a process known as tokenization. Effective tokenization defines the basic units of representation, directly influencing representation learning. We adopt Byte Pair Encoding (BPE) [13, 14] to construct vocabulary:

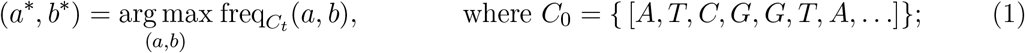

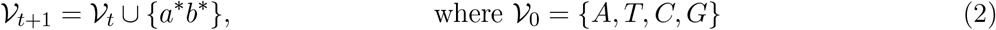

Here, *C*_*t*_ denotes the eccDNA corpus and 𝒱_*t*_ the vocabulary consisting of the nucleotide patterns at iteration *t*. The symbols *a* and *b* represent two adjacent nucleotide tokens in the corpus, and 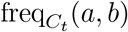 denotes their co-occurrence frequency. The BPE algorithm iteratively merges the most frequent adjacent pair of symbols (*a*^∗^, *b*^∗^), where the superscript ^∗^ indicates that the pair is selected as the most frequent in the current iteration *t*. The upper bound on *t* is determined by the vocabulary size. The merged token *a*^∗^*b*^∗^ is then added to the vocabulary. Fixed-length k-mer tokenization, as adopted in models such as DNABERT [20] and the Nucleotide Transformer [8], treats all substrings of length k equally and leads to exponential vocabulary growth as k increases. In contrast, modern genomic models such as DNABERT-2 employ BPE to alleviate this limitation [7]. Following this direction, we use BPE with a vocabulary size of 4096 to learn variable-length nucleotide patterns based on frequency, allowing the tokenizer to capture recurrent nucleotide patterns while keeping sequence length compact and enabling efficient modeling of ultra-long eccDNA.

### 2.2 Circular Data Augmentation

eccDNAs are circular molecules, yet standard linear representations ignore potential head–tail dependencies. Prior studies suggest the ends of eccDNA molecules often contain repetitive elements involved in circularization [21]. To preserve this circular topology, we introduce a data augmentation strategy that appends the first 64 tokens of each sequence to its end. Formally, for a sequence with *L* tokens, we construct the augmented input as 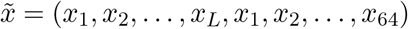. This lightweight operation preserves the continuity between the head and tail of the eccDNA sequence, enabling the model to capture long-range dependencies across the circular junction.

We choose 64 tokens because they provide sufficient contextual coverage for most circular junctions while also enabling efficient large-scale pretraining (see Appendix A.9 for more details). To achieve successful pretraining, we empirically chose to append approximately 25% of each sequence to its end. Most eccDNA sequences used for pretraining contain fewer than 256 tokens, so appending 25% corresponds to 64 tokens, aligning with our design choice. We also empirically validate the benefits of this augmentation in Section 3.2. Across all downstream tasks, integrating this circular augmentation leads to consistent performance improvements.

### 2.3 Bidirectional Mamba-2 Encoding

We aim to build a sequence encoder that can process eccDNA in both directions and remain aware of its circular structure while staying efficient enough to handle ultra-long inputs. To achieve this, we adopt Mamba-2 [15], a state-space model that processes sequences one token at a time instead of comparing all pairs of positions as in attention. This formulation enables linear-time computation and makes it feasible to scan megabase-scale DNA.

Formally, at each position *t*, the model maps the input token *x*_*t*_ to an embedding

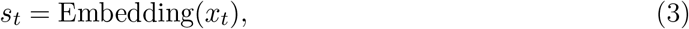

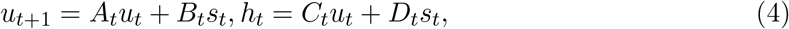

where *u*_*t*_ summarizes the information accumulated up to step *t*, and *h*_*t*_ is the contextual output for that position. These updates rely on fixed-cost operations, allowing the encoder to move through the sequence in a single pass, much like scanning along a chromosome. Meanwhile, the dynamically generated matrices *A*_*t*_, *B*_*t*_, *C*_*t*_, *D*_*t*_ introduce nonlinear mixing that enables the model to capture complex sequence patterns while preserving the efficiency of state-space recurrence.

eccDNAMamba uses two independent Mamba-2 encoders that scan the sequence in opposite directions, allowing the model to capture both long-range interactions and circular dependencies in ultra-long eccDNA while maintaining linear-time computation. The detailed mathematical formulation of Mamba-2 is provided in Appendix A.2.

As shown in Fig. 1, each eccDNA sequence is first tokenized into nucleotide pattern embeddings using the BPE vocabulary described in Section 2.1, which reduces redundancy and shortens the effective sequence length. The model then applies the circular augmentation described in Section 2.2, where the first 64 tokens are appended to the end of the sequence to explicitly expose the head–tail junction. The resulting augmented sequence 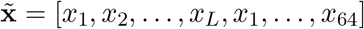 is fed into the forward and reverse Mamba-2 encoders.

The forward encoder scans the augmented sequence in its natural order, while the reverse encoder reads the reversed sequence:

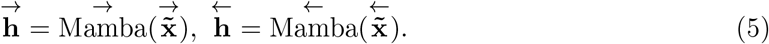

Each *h*_*i*_ represents the contextual embedding of token *x*_*i*_, capturing both local patterns and long-range dependencies. The two directional streams are then merged through a shared projection layer to form unified sequence representations:

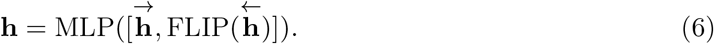

Here, the FLIP operator reverses the temporal order of the reverse encoder output, ensuring that forward and reverse positions are aligned, particularly at the head–tail junction. This alignment preserves symmetric contextual information and maintains awareness of the circular topology. After the large-scale pretraining phase, we further finetune the model on a specific biological task. This finetuning helps the model adapt its general understanding of eccDNA to the particular problem.

### 2.4 Pretraining

During pretraining, we further enhance representation learning by masking contiguous spans of tokens rather than isolated ones [22]. Each span consists of roughly three consecutive tokens and covers about 15% of the sequence in total. 80% of masked tokens are replaced with `[MASK]`, 10% are replaced with random tokens, and 10% remain unchanged. Formally, given a augmented sequence 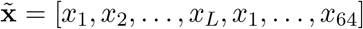 and a set of masked spans ℳ, the model predicts the original tokens by minimizing ℒ, the span-masked cross-entropy loss:

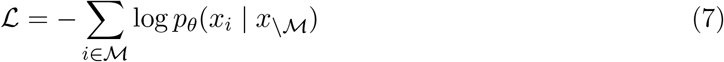

Compared to standard pretraining loss, our method jointly reconstructs tokens within a span, capturing intra-span dependencies and contextual coherence, properties crucial for long-range circular dependencies.

## 3 Experimental Setup

### 3.1 eccDNA Multi-Task Benchmark

To comprehensively assess model performance, we introduce the EccDNA Multi-Task Evaluation benchmark and assemble a unified eccDNA corpus by standardizing entries from CircleBase and eccDNAdb [5, 6], with additional details provided in Appendix A.1.

#### Cancer vs. healthy eccDNA discrimination

These tasks evaluate whether the sequence of an eccDNA molecule alone carries predictive signals associated with oncogenic activity. Cancer-derived eccDNAs are closely linked to oncogene amplification, transcriptional dysregulation, tumor evolution, and therapeutic resistance [1, 3, 4]. By asking whether cancer-associated eccDNAs can be distinguished from healthy eccDNAs using sequence alone, we test whether eccDNA sequences encode intrinsic molecular signatures that are informative even without orthogonal measurements such as expression. We consider two length regimes: short eccDNAs (<10 kbp) and ultra-long eccDNAs (10–200 kbp). This design separates baseline performance from the ability to model ultra-long eccDNAs.

#### EccDNA copy-number level prediction

In this task, the model is asked to infer from the eccDNA sequence alone whether the copy number is low or high. Because copy-number alterations are a major mechanism of oncogene amplification, we constructed this benchmark using human cancer eccDNAs from eccDNAdb [6], with segment-level genomic coordinates and copy-number annotations. Each eccDNA sequence was reconstructed from hg19 by extracting its annotated segments, reverse-complementing negative-strand segments, and concatenating fragments in the recorded order. Entries with missing copy-number values were excluded. We then binarized the remaining examples using two biologically motivated thresholds, copy number > 4.5 and copy number > 10, minimum threshold used to detect amplification event and median copy number of oncogene containing eccDNA defined in recent publications, respectively, to evaluate predictive signal across broader and more extreme amplification regimes [2, 17].

The dataset is consistently split with a train-to-test ratio of 80:20. To address potential data leakage, we evaluated sequence redundancy across the entire dataset using MMseqs2 clustering [23] at an 80% identity threshold. MMseqs2 computes this non-redundancy ratio by clustering sequences that meet the 80% identity cutoff and then taking the fraction of eccDNAs that do not fall into a cluster with a close homolog. Biologically, a higher ratio indicates a more diverse dataset with fewer near-duplicate or highly similar eccDNAs shared across splits, whereas a lower ratio indicates greater sequence redundancy and potential homology-driven leakage. As detailed in Table 1, the split datasets show high non-redundancy ratios (>0.95 under this threshold), suggesting low sequence overlap across the entire dataset. For each task, we add a classification head on top of the model and fine-tune only the classification head to perform the corresponding classification task. By standardizing this homology-aware sampling, along with task definitions and evaluation metrics, the eccDNA multi-task evaluation benchmark provides a robust framework for testing sequence modeling approaches.

**Table 1.** Overview of the two main tasks, including clustering statistics (percentage) for data redundancy reduction.

| Task | Sequence Length | Num. Train | Num. Eval | Non-redundancy Ratios | Dataset |
| --- | --- | --- | --- | --- | --- |
| Cancer vs. healthy eccDNA discrimination (short) | <10k | 16000 | 4000 | 97.04% | CircleBase [5] |
| Cancer vs. healthy eccDNA discrimination (ultra-long) | 10k–200k | 5011 | 811 | 95.66% | CircleBase [5] |
| EccDNA copy-number level prediction | <1M | 760 | 192 | 97.92% | eccDNAdb [6] |

### 3.2 Evaluation Metrics

We employed the standard cross-entropy loss to train all classification heads. Given an input augmented sequence 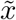, we first obtain unified sequence representations `h` using bidirectional Mamba-2 encoder (Sec. 2.3). We then apply masked mean pooling over `h` to derive a sequence-level representation *h*_pool_. The task-specific classification head *f*_*θ*_ then produces a predicted probability distribution 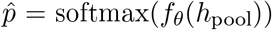 for the ground-truth label *y* ∈ {1, …, *C*}. The loss is computed as:

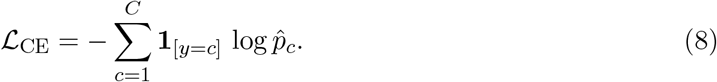

We use the Matthews Correlation Coefficient (MCC) as the evaluation metric. MCC takes into account true positives (*TP*), true negatives (*TN*), false positives (*FP*), and false negatives (*FN*), providing a measure of overall performance through

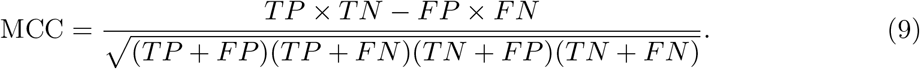

We also use F1 score and Area Under the Receiver Operating Characteristic curve (AUROC) to evaluate model predictions (Appendix A.5). The full experimental results are reported in Appendix A.6.

### 3.3 Baselines

We compare eccDNAMamba with three representative baselines for genomic sequence analysis covering both attention-based and state-space paradigms: DNABERT2 [20], a Transformer pretrained on genomic k-mers; HyenaDNA [9], a long-range implicit-convolution model that scales efficiently; Caduceus [11], a bidirectional state-space model with base-level tokenization that captures symmetric context.

In addition, we include eccDNAMamba–1M (w/o CA) as training eccDNAMamba without circular augmentation during both the pretraining and fine-tuning phases. This is an ablation model to verify the effect of circular augmentation.

Each model name is suffixed with its maximum supported input length, and any sequence exceeding the maximum length supported by a given baseline model is truncated during tokenization. All baselines are fine-tuned on the same eccDNA Multi-Task benchmark using their recommended learning rates, while eccDNAMamba and its ablation variant are fine-tuned with a learning rate of 3e-4. Each experiment is repeated with three random seeds, and we report the mean and standard deviation of all metrics (MCC, F1, and AUROC) for fair comparison.

## 4 Results

### eccDNAMamba identifies cancer-associated eccDNAs across short and ultra-long length regimes

We first evaluate the ability of models to identify cancer-associated eccDNAs across short and ultra-long length regimes. As shown in Figure 2, eccDNAMamba achieves the best overall performance across both regimes, reaching 59.0% MCC and 79.3% F1 on short sequences, and 57.9% MCC and 82.1% F1 on ultra-long ones. Its ablated variant without circular augmentation shows slight degradation, confirming the effectiveness and necessity of topology-aware modeling. Beyond oncogene-associated coding content, later interpretability analyses show that eccDNAMamba also assigns high attribution to non-coding regulatory regions and repeat-associated sequences, suggesting that the model is capturing broader features of cancer eccDNA architecture, including sequence elements linked to transcriptional dysregulation and genomic rearrangement, rather than relying solely on coding sequence identity (see Section 5).

**Figure 2:**
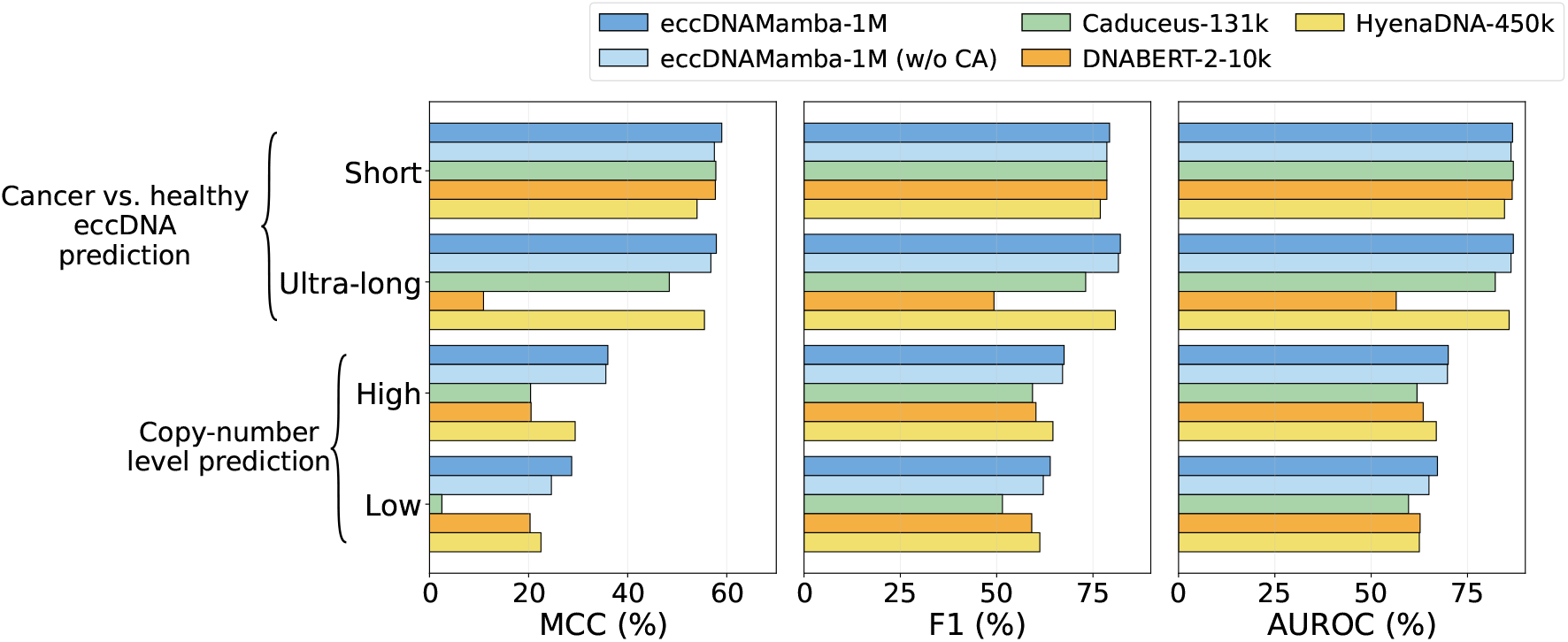
Performance of different models on **Cancer vs. healthy eccDNA prediction** and **eccDNA copy-number level prediction.** EccDNAMamba outperforms all baseline models on eccDNA sequences shorter than 10k (short) and between 10k and 200k (ultra-long). It also shows consistently robust performance under settings of the copy-number classification task.

In contrast, DNABERT-2 and Caduceus only slightly outperform HyenaDNA on short sequences with MCC values of 57.7% and 57.8% compared to 54.0%, but their performance drops sharply on long sequences. HyenaDNA remains relatively robust with 55.5% MCC, whereas DNABERT-2 collapses to 10.9%. DNABERT-2 is inapplicable to sequences longer than 10k and requires truncation for ultra-long sequences. This highlights the importance of modeling long-range dependencies for maintaining the topological integrity of eccDNA.

### eccDNAMamba infers eccDNA copy-number levels

eccDNA is routinely replicated in cancer cells, selectively amplifying sequences that can exacerbate the cells’ oncogenic potential [2]. Conventional copy-number estimation depends on costly high-coverage sequencing or hybridization-based assays, rendering it less practical or scalable than sequence-only analysis. To address these issues, we evaluate whether eccDNA copy-number levels can be inferred directly from sequences. To make this task comparable and robust under a limited sample size, we evaluate two thresholded label settings: a low threshold (>4.5) for broader copy-number elevation and a high threshold (>10) for more extreme amplification.

As shown in Figure 2, under a small-sample setting of only 760 training sequences (Table 1), Caduceus and DNABERT-2 show substantial degradation, with MCC values of 20.4% and 20.5% at the higher threshold and 2.5% and 20.3% at the lower threshold. HyenaDNA performs more stably with 29.4% MCC and 64.6% F1. In contrast, eccDNAMamba achieves the best and most stable results across both threshold settings, reaching 36.0% MCC and 67.5% F1 at the higher threshold and 28.7% MCC and 63.9% F1 at the lower threshold, effectively capturing copy-number–related signals despite limited data. To the best of our knowledge, we demonstrated for the first time that copy number variations can be effectively predicted from sequence information alone, highlighting the potential utility of sequence-based models for scalable copy-number screening.

### eccDNAMamba enables memory-efficient fine-tuning

we visualize the memory footage of different models during the fine-tuning process in Figure 3. During fine-tuning, eccDNAMamba uses only 40% memory, which is 30% lower than HyenaDNA and Caduceus, and 50% lower than DNABERT2. The near-constant footprint for eccDNAMamba aligns with our analysis (Appendix A.4), in contrast to attention-based baselines whose usage grows with sequence length and runtime.

**Figure 3:**
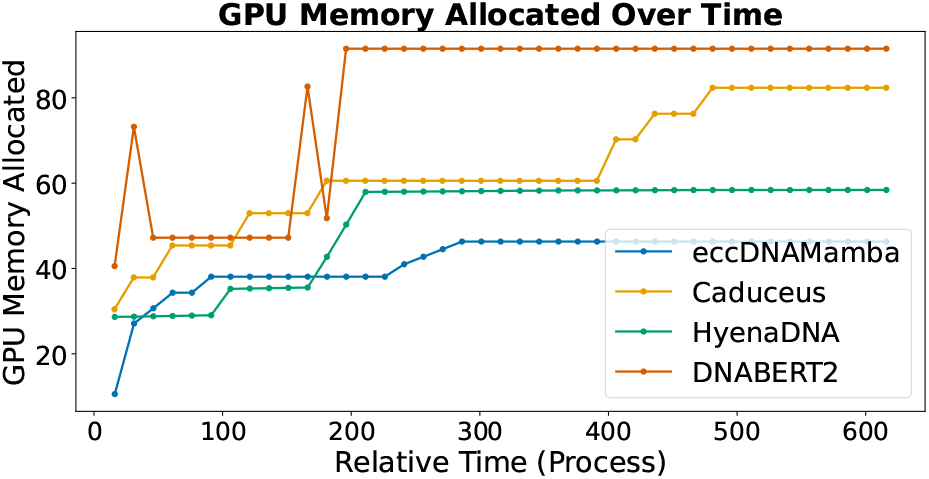
GPU memory usage over time for eccDNAMamba, Caduceus, HyenaDNA, and DNABERT-2, showing eccDNAMamba’s lower and stable memory footprint

## 5 Biological Interpretations

To understand what drives eccDNAMamba predictions, we use Integrated Gradients (IG) for analysis, because it satisfies completeness and measures each token’s true contribution by integrating gradients along a path, avoiding the noise of single-step saliency, while remaining naturally compatible with Mamba’s attention-free, ultra-long-sequence architecture. IG assigns token-level attributions along a path from a pad baseline *x*_*pad*_, which is fully filled with PAD tokens, to the input sequence *x*. For each token *x*_*i*_, IG integrates the gradient of the model’s output with respect to *x*_*i*_ along the straight-line path *x*_*α*_ = *x*_*pad*_ + *α*(*x* − *x*_*pad*_), *α* ∈ [0, 1], using the logit difference between the target and other classes as the decision signal:

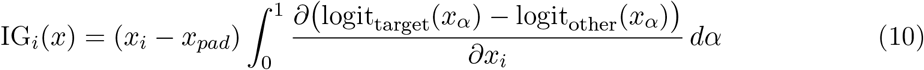

Therefore, IG yields a principled, faithful importance score for each nucleotide, capturing sensitivity across the entire interpolation rather than a single gradient snapshot, and manages to reveal sequence features that drive eccDNAMamba’s decisions.

### IG reveals distinct landscapes across prediction groups

After stratifying sequences by prediction outcome, markedly different IG profiles emerged between eccDNA sequences predicted as cancerous and those predicted as healthy. In the ultra-long model (10-200k), cancerous sequences (true positives) exhibited consistent positive signal throughout the entire sequences, with slightly stronger values in the first half. In contrast, healthy sequences (true negatives) present a prominent bipartite IG profile with negative signals at the first half of the sequence and positive signals at the second half (Figure 4). These differences highlight the global differences our model is able to clearly distinguish between healthy and cancerous eccDNA sequences.

**Figure 4:**
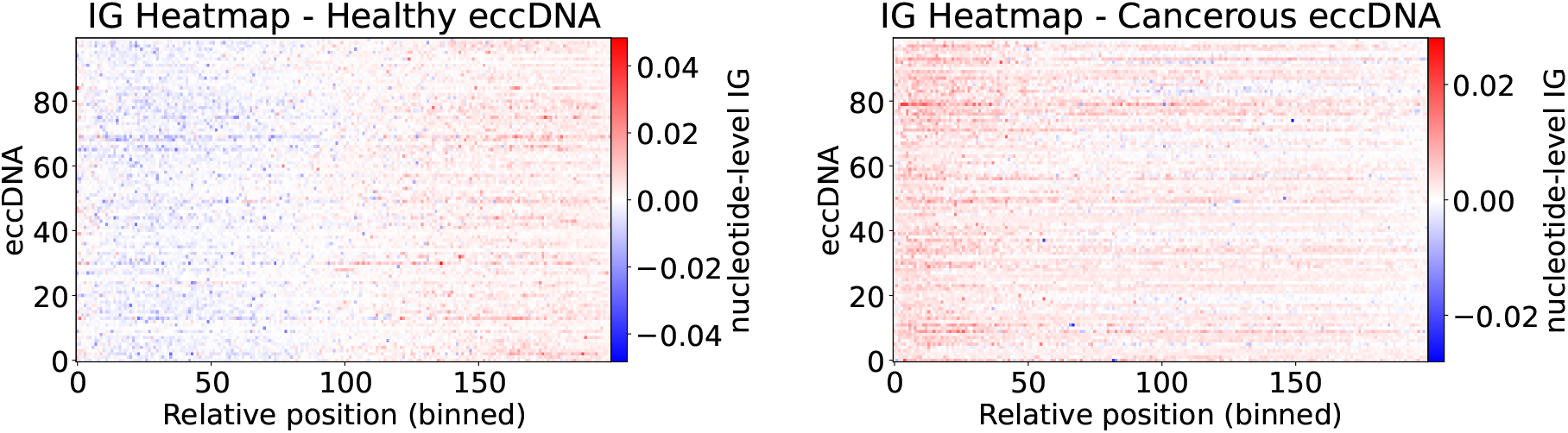
IG values across eccDNAs of correctly predicted healthy (left) and cancerous (right) sequences by the ultra-long model.

### Regulatory element groups and transposable element families receive differential attribution

Using ENCODE annotations (ENCODE cCRE v4) [24], we analyzed the aggregated IG contributions of each class of regulatory elements. Across both short (<10 kbp) and ultra-long (10–200 kbp) settings, cCRE categories such as PLS (promoters), pELS (proximal enhancers), and dELS (distal enhancers) contribute significantly higher per-base |IG| values than length- and context-matched background sequences randomly selected from other regions of the same eccDNAs (Figure 5A,B). Based on RepeatMasker annotations [25], we also observed retrotransposon (RTE) families-specific |IG| patterns: Alu elements showed significantly lower, while LINE-1 and ERV families exhibited higher per-base |IG| than matched background (Figure 5C). These results indicate that eccDNAMamba systematically prioritizes certain regulatory DNA and selectively attends to specific TE classes, consistent with eccDNA’s ability to harbor RTE and promoter/enhancer-like sequences that can amplify oncogenic programs [1, 26].

**Figure 5:**
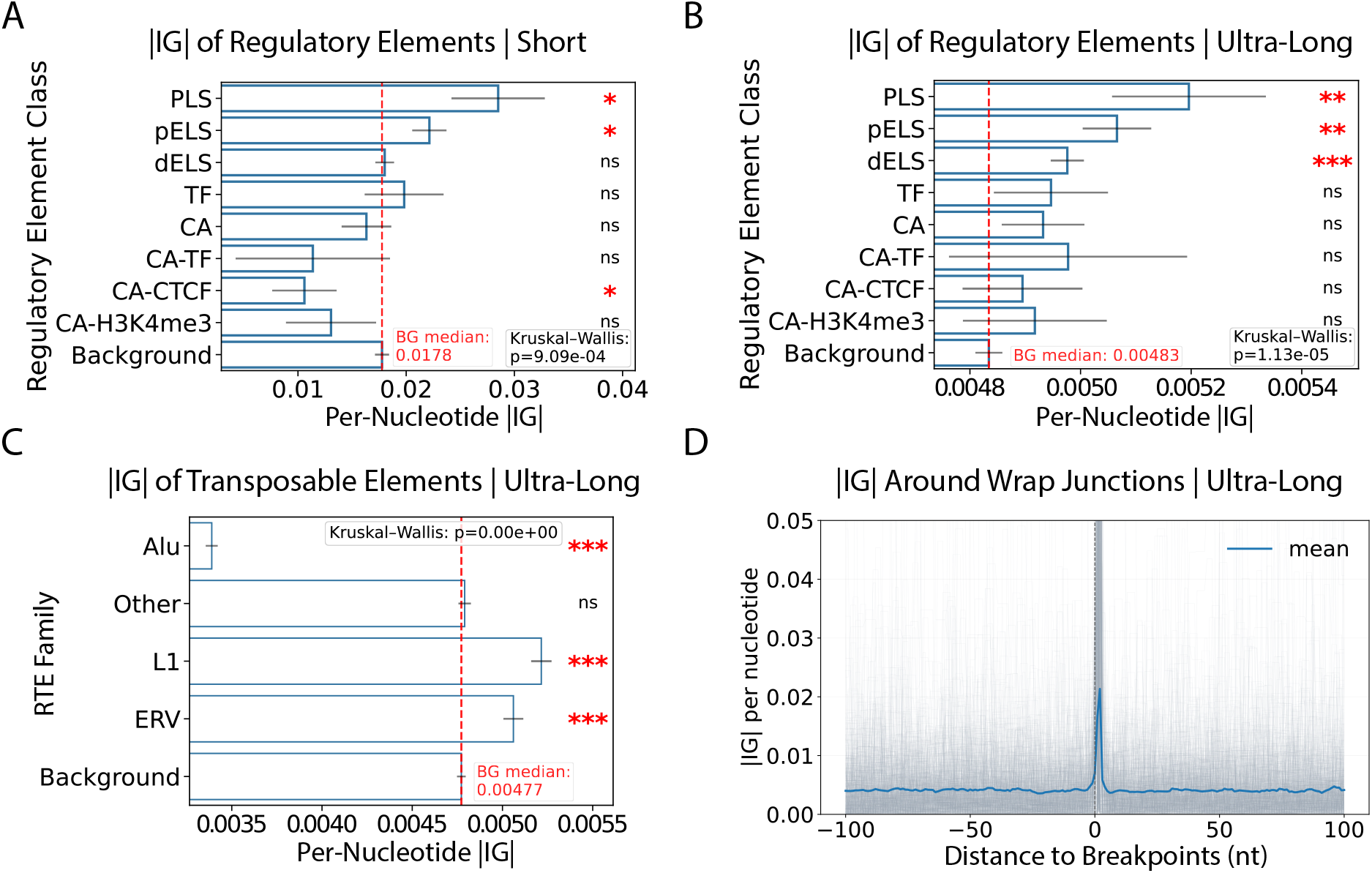
(A,B) Per-nucleotide |IG| values across gene regulatory elements from the short (below10k) model (A) and ultra-long (10-200k) model (B). (C) Per-nucleotide |IG| values across transposable elements from the ultra-long (10-200k) model. (D) |IG| peaks at breakpoints of eccDNA by the ultra-long model. Kruskal Wallis test determines significant differences between groups. Stars indicate significance from two-tailed Mann–Whitney U test, as compared to the “Background.” (“*” = p < 0.05, “**” = p < 0.01, “***” = p < 0.001)

### Breakpoint-centered attribution highlights circular topology

Aggregating IG scores around the breakpoints revealed a pronounced enrichment at the head–tail junction of sequences (Figure 5D). This enrichment validates that the circular augmentation strategy is recognized and important to our model, capturing head–tail dependencies. It provides direct interpretability evidence on the role of augmentation in the model’s fidelity to eccDNA topology.

### eccDNAMamba discovers known and novel motifs relevant for cancer eccDNA

Interpreting high–|IG| subsequences from eccDNAMamba using XSTREME [27] uncovered 69 motifs, 40 of which (58%) matched known matrices by TomTom(CIS-BP_2.00 | *Homo sapiens*) [28, 29]. Twenty-three motifs were significantly enriched in cancer eccDNA (log_2_ OR > 0; *q* < 0.05), led by A/T-rich seeds such as **TAATTAATAT** (log_2_ OR = 15.65; *q* = 9.0 × 10^−12^). Among mapped examples, **TTTCCTGAGAAAT** aligns to a STAT-like matrix (M08173_2.00), **GTAAACAATAAACA** aligns to a FOX-like matrix (M03016_2.00), and **TAATTAATAT** aligns to an ARID-family matrix (M01659_2.00), collectively pointing to transcription-factor programs frequently implicated in oncogenesis (cytokine/STAT signaling, FOX pioneer-factor networks, and A/T-binding ARID factors) [30, 31, 32]. Strikingly, only 8/23 motifs aligned to known motifs, all of which match transcription factor families relevant to cancer biology. This indicates that the other 15/23 cancer-enriched motifs, which lack an unambiguous database match, could serve as a resource to reveal previously unannotated sequence patterns associated with cancer eccDNA. These results could provide mechanistic interpretability and testable hypotheses about regulatory grammar on cancer-derived eccDNA.(Figure 6 and Appendix A.10)

**Figure 6:**
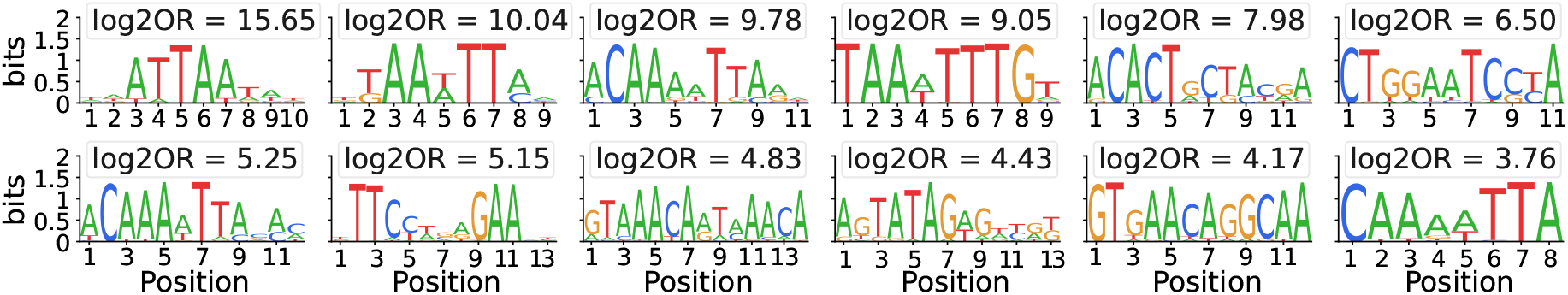
Representative cancer-overrepresented motifs discovered by IG analysis of eccDNAMamba.

Taken together, these analyses provide three insights. eccDNAMamba learns to prioritize biologically important elements and motifs across length regimes. The model distinguishes correct from incorrect predictions through differences in IG landscapes. Our circular augmentation is beneficial for state-of-the-art performance and interpretable: it induces consistent, localized attribution signals at the head–tail junction, reflecting the circular topology of eccDNA.

## 6 Discussion

eccDNAMamba presents a topology-aware, bidirectional state-space framework that efficiently models the circular and ultra-long characteristics of eccDNA. By integrating bidirectional Mamba-2 encoding, motif-level BPE tokenization, and lightweight head–tail augmentation, the model captures wrap-around dependencies and long-range interactions in linear time. Integrated Gradients analyses further confirm the biological interpretability of the model. The breakpoint-centered attribution analysis indicates that eccDNAMamba is sensitive to junction-proximal sequence context, motivating future work to test whether these signals correspond to known biogenesis-related features such as repeats or microhomology.

Despite its strong performance, sequence-only models cannot capture all determinants of eccDNA behavior, particularly epigenetic state and cell-type-specific regulation. Our results therefore establish sequence as an highly-informative modality, not a complete mechanistic description. Additionally, current training data are primarily human-centric, limiting cross-species generalization. Future work will expand pretraining to broader multi-species eccDNA corpora and apply the framework to more diverse downstream tasks.

## A Supplementary Information

### A.1 Dataset

As summarized in Table 2, we assemble a unified eccDNA corpus by standardizing several public resources: CircleBase [5], which provides 601,036 human eccDNAs including both healthy and cancer-associated sequences; eccDNAdb [6], which contributes 1,270 eccDNAs identified from 480 samples spanning 42 cancer types, derived from a total of 3,395 tumor samples covering 57 cancers across patient tissues and cancer cell lines; and eccDNA Atlas [16], which aggregates 629,987 eccDNAs across multiple species, expanding beyond human to include *Mus musculus, Saccharomyces cerevisiae, Gallus gallus*, and *Arabidopsis thaliana*.

**Table 2.** List of eccDNA datasets used in this study.

| Dataset | Species | Number of eccDNAs | Additional Information |
| --- | --- | --- | --- |
| CircleBase [5] | *Homo sapiens* | 601,036 | Healthy and cancer-associated eccDNAs |
| eccDNA Atlas [16] | Multi-species | 629,987 | Multi-species eccDNAs |
| eccDNAdb [6] | *Homo sapiens* | 1,270 | eccDNAs from Cancer samples and cell lines |

### A.2 Mamba-2 Architecture

The Mamba-2 [15] is a linear state-space model (SSM) that parameterizes sequence dynamics as a continuous-time linear system:

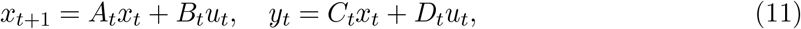

where *u*_*t*_ ∈ ℝ^*d*^ is the input token embedding, *x*_*t*_ is the latent state, and *y*_*t*_ is the output at time *t*. This formulation enables constant-time updates per token and thus overall 𝒪 (*L*) complexity for a sequence of length *L*, in contrast to 𝒪 (*L*^2^) attention. The parameters (*A*_*t*_, *B*_*t*_, *C*_*t*_, *D*_*t*_) are dynamically generated via depthwise convolutions and gated activations, allowing nonlinear feature mixing while maintaining linear-time recurrence.

### A.3 Pretraining

We adopt a hybrid method combining contextual embedding-level control and masking as reported in the previous section during pretraining.

We pre-train the model using a span-masked language modeling (SpanMLM) objective adapted for eccDNA modeling. Specifically, we use a bidirectional variant of the Mamba architecture, denoted as **BiMambaForMaskedLM**, initialized from the state-spaces/mamba2-130m configuration. The model is trained from scratch on a custom eccDNA corpus, tokenized using a byte-pair encoding vocabulary. The tokenizer includes the special tokens `[PAD]`, `[UNK]`, `[MASK]`, and `[CLS]`.

Let the input sequence be denoted as

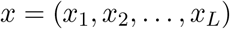

and let ℳ ⊂ {1, …, *L*} be a set of span-masked positions. The model is trained to reconstruct the original tokens at these positions by minimizing the span-masked cross-entropy loss:

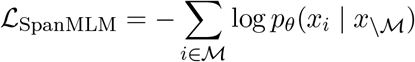

where *p*_*θ*_ denotes the model’s conditional output distribution and *x*\_ℳ_denotes the input with masked spans.

Training is conducted using the HuggingFace `Trainer` API. We set the effective batch size to 144 by using 6 samples per GPU and accumulating gradients over 8 steps. The model is trained for 3 epochs using a learning rate of 5 × 10^−4^, with a linear warmup over 6% of total steps. We use the AdamW optimizer with a weight decay of 0.01. All training runs use BF16 mixed precision and are executed on a single node with 3 NVIDIA L40S GPUs.

As shown in Table 3,we filter out sequences longer than 10kbp, resulting in a final corpus of 1, 087, 886 eccDNA sequences totaling 524 million base pairs on the dataset we introduced in Appendix A.1. After applying BPE tokenization, this corpus comprises approximately 101.5 million tokens, yielding an average token length of 5.16 base pairs per token after BPE tokenization.

**Table 3.** Statistics of the eccDNA corpus after filtering (sequences longer than 10kbp removed) and BPE tokenization.

| Number of Sequences | Total Base Pairs | Tokens (BPE) | Average Token Length (bp) |
| --- | --- | --- | --- |
| 1,087,886 | 524M | 101.5M | 5.16 |

**Table 4.** Performance of different models on Cancer vs. healthy eccDNA prediction.

| Task | Model | MCC (%) | F1 (%) | AUROC (%) |
| --- | --- | --- | --- | --- |
| Cancer vs. healthy eccDNA discrimination (short) | eccDNAMamba-1M | **59.0±0.6** | **79.3±0.2** | **86.7±0.3** |
|  | eccDNAMamba-1M (w/o CA) | 57.5±1.6 | 78.6±0.7 | 86.3±0.1 |
|  | Caduceus-131k | 57.8±1.2 | 78.6±0.5 | 86.9±0.5 |
|  | DNABERT-2-10k | 57.7±1.3 | 78.6±0.6 | 86.6±0.5 |
|  | HyenaDNA-450k | 54.0±0.5 | 76.9±0.2 | 84.6±0.2 |
| Cancer vs. healthy eccDNA discrimination (ultra-long) | eccDNAMamba-1M | **57.9±4.5** | **82.1±1.1** | **86.9±0.8** |
|  | eccDNAMamba-1M (w/o CA) | 56.8±4.3 | 81.6±1.0 | 86.3±1.2 |
|  | Caduceus-131k | 48.4±3.5 | 73.1±2.6 | 82.2±3.3 |
|  | DNABERT-2-10k | 10.9±3.8 | 49.3±6.7 | 56.5±2.0 |
|  | HyenaDNA-450k | 55.5±2.1 | 80.8±2.0 | 85.8±1.7 |

**Table 5.** Performance of different models on eccDNA copy-number level prediction.

| Task | Model | MCC (%) | F1 (%) | AUROC (%) |
| --- | --- | --- | --- | --- |
| eccDNA copy-number level prediction (copy number > 10) | eccDNAMamba-1M | **36.0±10.6** | **67.5±5.5** | **70.0±4.6** |
|  | eccDNAMamba-1M (w/o CA) | 35.6±8.0 | 67.1±4.4 | 69.8±5.4 |
|  | Caduceus-131k | 20.4±22.8 | 59.3±10.4 | 61.9±6.8 |
|  | DNABERT-2-10k | 20.5±11.3 | 60.2±5.5 | 63.5±7.8 |
|  | HyenaDNA-450k | 29.4±8.9 | 64.6±4.4 | 66.9±5.8 |
| eccDNA copy-number level prediction (copy number > 4.5) | eccDNAMamba-1M | **28.7±4.5** | **63.9±1.9** | **67.2±2.3** |
|  | eccDNAMamba-1M (w/o CA) | 24.6±6.2 | 62.1±2.9 | 65.0±0.6 |
|  | Caduceus-131k | 2.5±12.5 | 51.5±5.8 | 59.7±4.5 |
|  | DNABERT-2-10k | 20.3±2.0 | 59.1±0.9 | 62.7±2.9 |
|  | HyenaDNA-450k | 22.5±17.6 | 61.2±8.7 | 62.5±9.4 |

### A.4 Efficiency

Let *s*_1:*L*_ ∈ ℝ^*d*^ be a single eccDNA sequence tokenized with byte-pair encoding (BPE), where *L* is the number of *tokens* and *d* the channel dimension. An eccDNAMamba block processes *u*_1:*L*_ as follows: (1) append a fixed *k*-token prefix to expose head–tail context, yielding length *L*′ = *L* + *k*; (2) run one forward scan and one reverse scan over the augmented sequence; (3) fuse the two output streams with a constant-cost operation (e.g., sum or concat+linear). We treat *k* as a small constant (default *k*=64), so *L*′ = Θ(*L*).

Each scan applies the per-step update

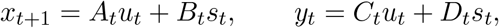

where computing (*A*_*t*_, *B*_*t*_, *C*_*t*_, *D*_*t*_) and performing the update at each position both cost *O*(1) work independent of *L*. Thus, a left-to-right pass costs *O*(*L*′), the reverse pass costs *O*(*L*′), and fusion is *O*(*L*′) in total.

Overall, the block runs in *O*(*L*′) + *O*(*L*′) + *O*(*L*′) = *O*(*L*) time; the bidirectional constant factor does not affect the asymptotics. Activation storage is *O*(*L*), and with standard chunking or rematerialization, the peak memory can be held to *O*(*B*) for a chosen block size *B* while preserving *O*(*L*) total work. Compared to self-attention’s *O*(*L*^2^*d*) cost, eccDNAMamba scales linearly in *L* and accommodates ultra-long eccDNA.

### A.5 Evaluation Metrics

We further report the F1 score, which balances precision and recall by taking their harmonic mean:

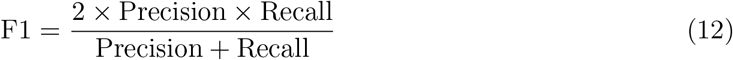

where precision and recall are defined as:

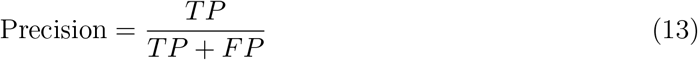

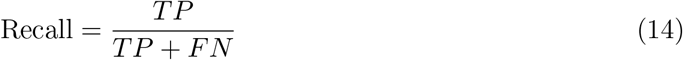

We also report the Receiver Operating Characteristic Area Under the Curve (AUROC), which measures the model’s ability to distinguish between positive and negative classes. It is computed by integrating the true positive rate (TPR) over the false positive rate (FPR) across all classification thresholds:

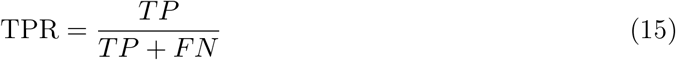

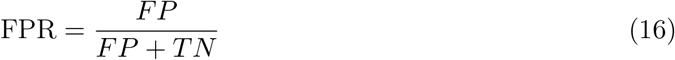

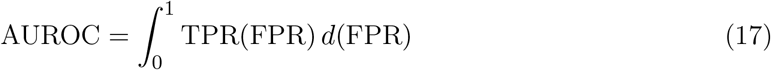

A higher AUROC indicates better discrimination ability of the model across varying thresholds.

### A.6 Full Experimental Results

We report the full experimental results for cancer-versus-healthy eccDNA discrimination and eccDNA copy-number level prediction in the following tables.

### A.7 Generalization on different species

We validate whether models can distinguish true eccDNA sequences from pseudo-eccDNA sequences that are randomly extracted from linear genomic regions of *Homo sapiens*. The pseudo-eccDNA sequences are generated to match the length distribution of true eccDNAs, ensuring that models cannot rely on length cues for classification. This task evaluates how well the model captures sequence features or compositions that distinguish true eccDNAs from random sequences.

As in Table 6, eccDNAMamba achieves the highest performance among all models with 42.7% MCC, 71.0% F1, and 75.8% AUROC, demonstrating superior ability to capture structural and contextual signals embedded in ultra-long eccDNAs. In contrast, DNABERT-2 and DeepCircle perform substantially worse with MCC values of 37.3% and 29.9%. Caduceus and HyenaDNA remain relatively robust yet still fall short of eccDNAMamba. Even though the mean MCC difference is 1.8 points (42.7 vs. 40.9), eccDNAMamba has tighter variance(± 1.8 vs. ± 3.9), indicating that its performance is more reliable, more reproducible, and less sensitive to random initialization. These results suggest that eccDNAMamba captures the fundamental biological features of real eccDNA, rather than relying on non-biological artifacts during classification. We also assess the generalization ability of our model by extending this task to *Gallus gallus* and *Arabidopsis thaliana* and observed similar trends.

**Table 6.**
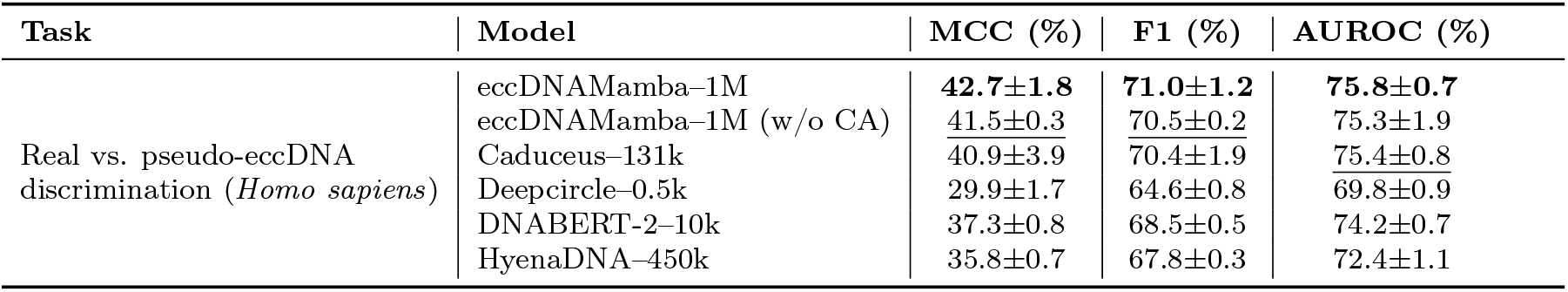
Performance of different models on Real vs. pseudo-eccDNA discrimination (*Homo sapiens*). eccDNAMamba achieves the best performance among all models, with a slight performance drop in its non-augmented version, yet still outperforms the baselines.

To evaluate the cross-species generalization ability of different genomic models, we further extended the real vs. pseudo-eccDNA discrimination task to two non-human species, *Arabidopsis thaliana* and *Gallus gallus*. This task examines whether a model trained primarily on human eccDNA can reliably identify genuine eccDNA structures in evolutionarily distant genomes. High performance on these datasets indicates robust representation learning and strong transferability beyond the training species.

As shown in Table 7, the performance of various genomic models on the *Arabidopsis thaliana* dataset. Across models, eccDNAMamba achieves the best MCC and F1 scores, while remaining competitive in AUROC. In contrast, Deepcircle and HyenaDNA exhibit substantial drops in MCC, suggesting that models designed for linear or short DNA contexts struggle to generalize to the *Arabidopsis thaliana* dataset.

**Table 7.** Results on *Arabidopsis thaliana*.

| Task | Model | MCC (%) | F1 (%) | AUROC (%) |
| --- | --- | --- | --- | --- |
| Real vs. pseudo-eccDNA discrimination (*Arabidopsis Thaliana*) | eccDNAMamba-1M | 49.7±1.0 | 74.5±0.7 | 79.2±3.1 |
| | eccDNAMamba-1M (w/o CA) | 48.1±3.7 | 73.7±1.7 | 78.8±3.1 |
| | Caduceus-131k | 43.6±5.3 | 71.2±2.6 | 79.6±4.6 |
| | Deepcircle-0.5k | 39.6±8.9 | 65.1±3.9 | 70.4±5.0 |
| | DNABERT-2-10k | 46.3±2.4 | 72.5±0.9 | 75.7±2.6 |
| | HyenaDNA-450k | 12.3±2.8 | 55.9±1.2 | 59.2±2.9 |

As shown in Table 8, eccDNAMamba achieves competitive performance with an MCC of 51.6%, closely matching Caduceus and its own ablated variant. This consistency highlights the robustness of eccDNAMamba in the *Gallus gallus* dataset. In contrast, transformer-based models such as DNABERT-2 and convolutional models like Deepcircle perform notably worse, reflecting limitations in generalizing to the *Gallus gallus* dataset.

**Table 8.** Results on *Gallus gallus*.

| Task | Model | MCC (%) | F1 (%) | AUROC (%) |
| --- | --- | --- | --- | --- |
| Real vs. pseudo-eccDNA discrimination (*Gallus Gallus*) | eccDNAMamba-1M | 51.6±4.2 | 75.1±2.5 | 81.6±2.4 |
| | eccDNAMamba-1M (w/o CA) | 50.4±2.1 | 75.0±1.2 | 81.5±1.6 |
| | Caduceus-131k | 52.1±0.8 | 76.0±0.4 | 82.8±0.1 |
| | Deepcircle-0.5k | 29.9±1.7 | 64.6±0.8 | 69.8±0.9 |
| | DNABERT-2-10k | 46.1±2.8 | 72.8±1.4 | 79.1±0.9 |
| | HyenaDNA-450k | 35.4±2.6 | 67.5±1.3 | 73.0±2.0 |

### A.8 Performance of Different Models on Cancer vs. Healthy eccDNA Discrimination Task (200k–1M)

In the ultra-long eccDNA regime (>200 kb), we observed a pronounced class imbalance where cancer-associated sequences overwhelmingly dominate, while healthy counterparts are sparsely represented (Figure 7). This skewed distribution creates a trade-off for the model: on one hand, sequence length itself becomes a strong implicit signal, encouraging the model to associate longer sequences with cancer; on the other hand, mitigating this bias risks diminishing sensitivity to genuine biological correlations between length and oncogenic features. Consequently, as shown in Table 9 eccDNAMamba maintains a high F1 score but exhibits a near-zero MCC, indicating a tendency to classify most ultra-long sequences as cancer-associated. This outcome highlights the need for future work to disentangle biological length effects from dataset-induced bias.

**Table 9.** Performance of different models on **Cancer vs. healthy eccDNA discrimination (200k–1M)**.

| Task | Model | MCC (%) | F1 (%) | AUROC (%) |
| --- | --- | --- | --- | --- |
| Cancer vs. healthy eccDNA discrimination (200k - 1M) | eccDNAMamba-1M | 0.0±0.0 | 88.1±0.7 | 55.8±0.5 |
| | eccDNAMamba-1M (w/o CA) | 0.0±0.0 | 88.1±0.7 | 53.3±1.8 |
| | Caduceus-131k | 0.0±0.0 | 88.1±0.7 | 61.6±2.0 |
| | DNABERT-2-10k | 10.7±1.8 | 65.6±9.5 | 61.2±1.5 |
| | HyenaDNA-450k | 5.1±4.5 | 86.4±2.9 | 68.8±7.7 |

**Figure 7:**
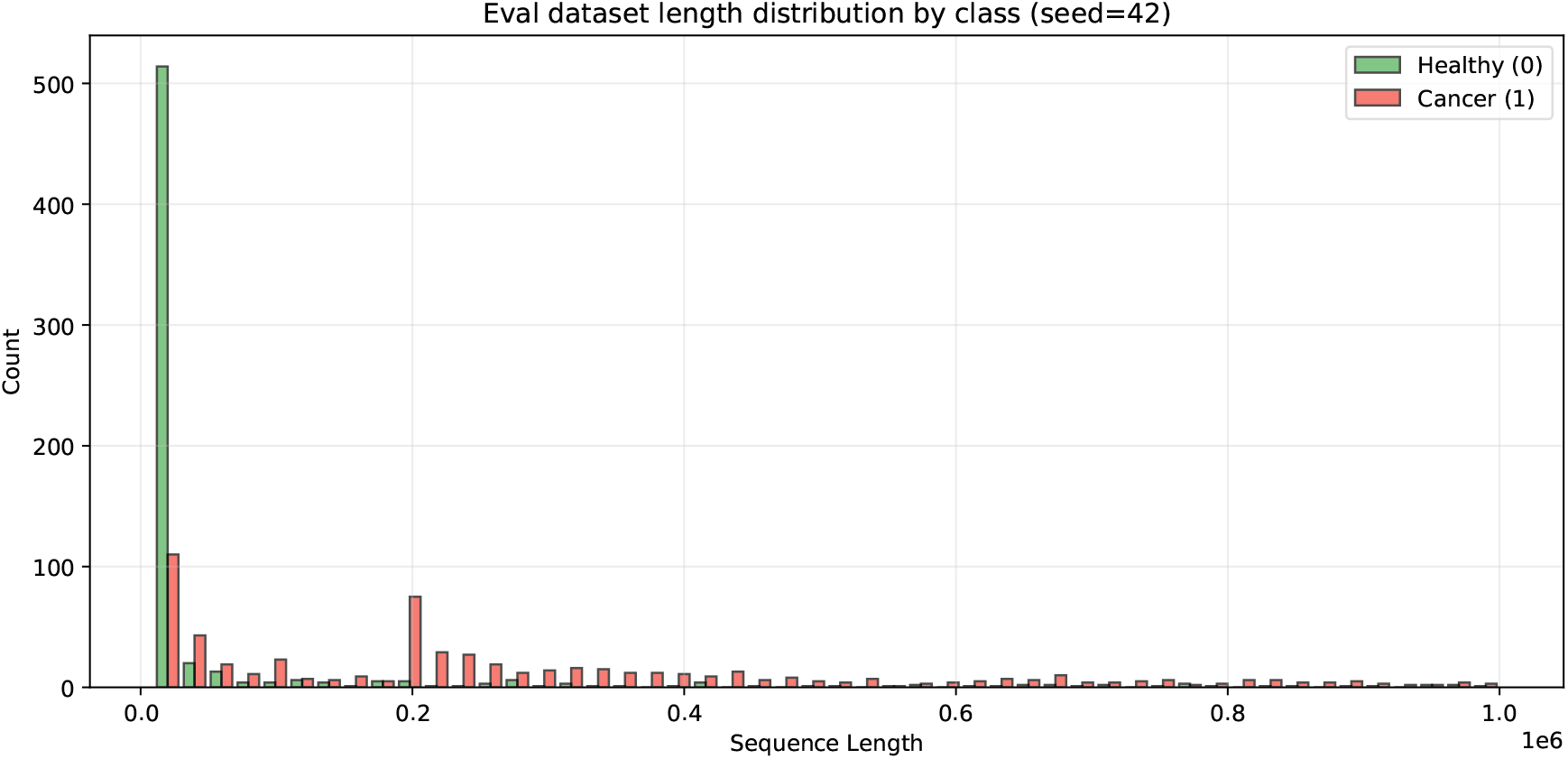
Length distribution of the evaluation dataset by class on **cancer vs. healthy eccDNA discrimination (200k–1M)**.

### A.9 Tokenization Statistics and Motivation for Circular Augmentation

We empirically chose to append approximately 25% of the sequence to its end to ensure successful pretraining. As shown in Figure 8a, most eccDNA sequences used for pretraining contain fewer than 256 tokens, so appending 25% corresponds to 64 tokens.

**Figure 8:**
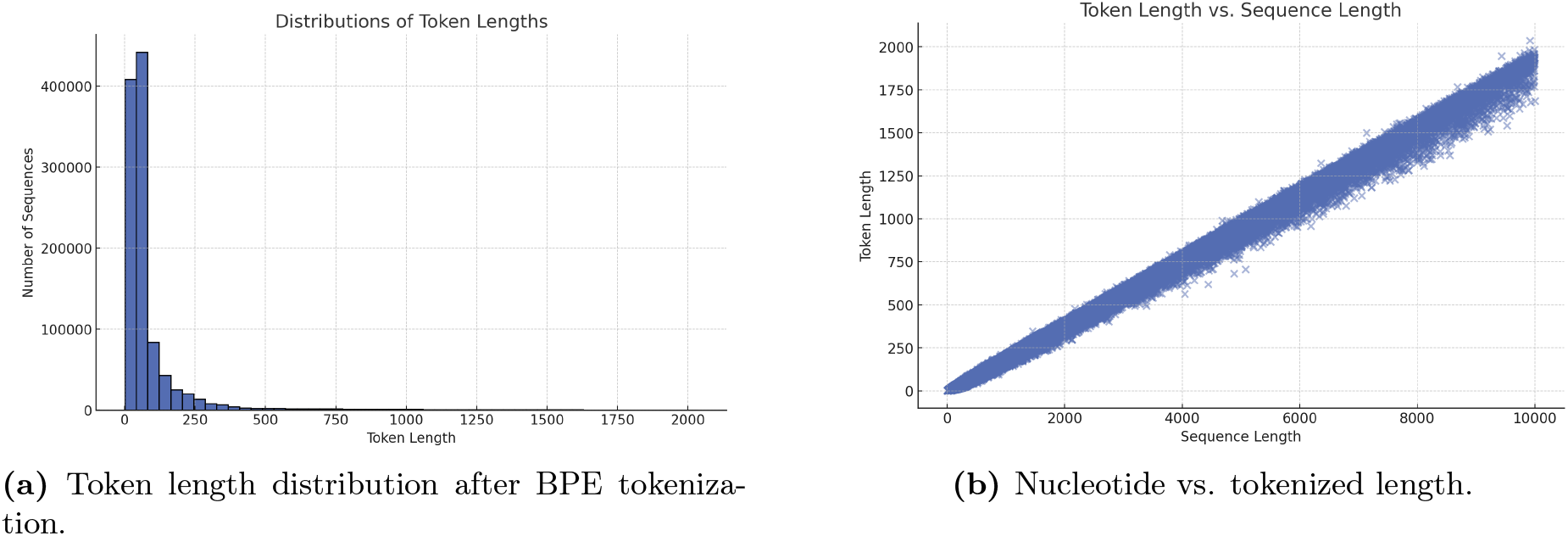
(a) Distribution of token lengths after BPE tokenization. Most sequences are tokenized into fewer than 256 tokens, with a long-tailed distribution extending to over 1000 tokens. (b) Relationship between nucleotide sequence length (in base pairs) and tokenized sequence length.

### A.10 Motif Identification

A full catalog of motifs enriched in cancer-derived or healthy eccDNA sequences, identified through our IG-based motif discovery pipeline, is shown in Figure 9. Of these, 8 of 23 cancer-enriched motifs and 1 of 6 healthy-enriched motifs matched known transcription factor motifs in CIS-BP v2. Notably, healthy-enriched motifs exhibit higher CG content, even though the underlying eccDNA sequences from cancer and healthy samples display similar overall CG composition (Table 10).

**Table 10.** Nucleotide composition across all sequences used for IG analysis and motif discovery. (n=726)

| Group | %A | %C | %G | %T | %GC |
| --- | --- | --- | --- | --- | --- |
| Cancer | 29.004 | 21.061 | 21.051 | 28.884 | 42.112 |
| Healthy | 28.137 | 21.848 | 21.775 | 28.239 | 43.623 |

**Figure 9:**
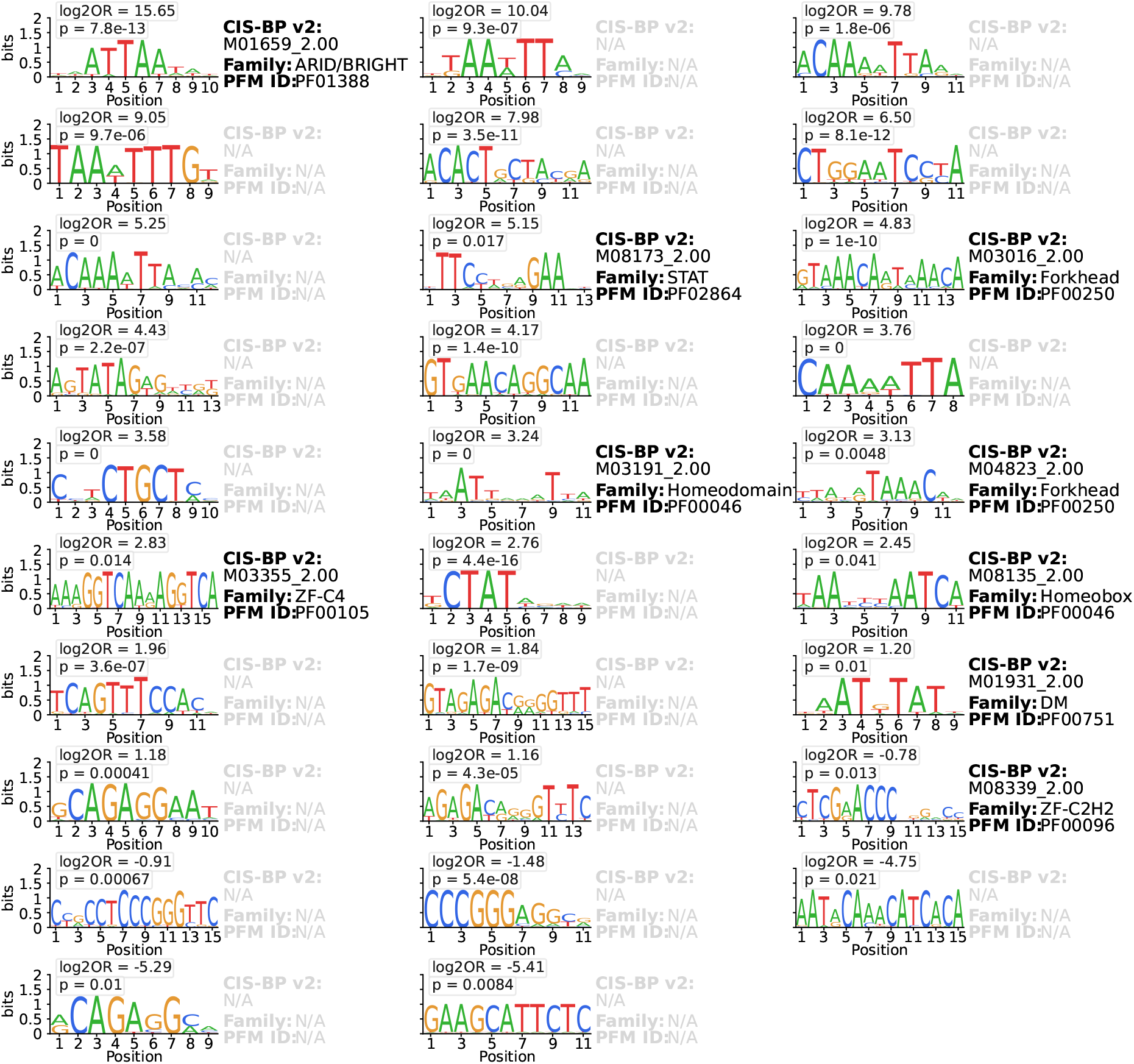
Motifs that are significantly over-represented in either cancer or healthy eccDNA sequences. Motifs with known matching in CIS-BP v2 are labeled with their CIS-BP motif ID, protein family, and Pfam ID.

## References

[1] Dong, Y. et al. Extrachromosomal dna (ecdna) in cancer: mechanisms, functions, and clinical implications. Frontiers in oncology 13, 1194405 (2023).

[2] Bailey, C., Pich, O., Thol, K. et al. Origins and impact of extrachromosomal dna. Nature 635, 193–200 (2024).

[3] Turner, K. M. et al. Extrachromosomal oncogene amplification drives tumour evolution and genetic heterogeneity. Nature 543, 122–125 (2017).

[4] Koche, R. P. et al. Extrachromosomal circular dna drives oncogenic genome remodeling in neuroblastoma. Nature genetics 52, 29–34 (2020).

[5] Zhao, X. et al. Circlebase: an integrated resource and analysis platform for human eccdnas. Nucleic acids research 50, D72–D82 (2022).

[6] Peng, L., Zhou, N., Zhang, C.-Y., Li, G.-C. & Yuan, X.-Q. eccdnadb: a database of extrachromosomal circular dna profiles in human cancers. Oncogene 41, 2696–2705 (2022).

[7] Zhou, Z. et al. Dnabert-2: Efficient foundation model and benchmark for multi-species genome. arXiv preprint arXiv:2306.15006 (2023).

[8] Dalla-Torre, H. et al. Nucleotide transformer: building and evaluating robust foundation models for human genomics. Nature Methods 22, 287–297 (2025). URL 10.1038/s41592-024-02523-z.

[9] Nguyen, E. et al. Hyenadna: Long-range genomic sequence modeling at single nucleotide resolution (2023). URL https://arxiv.org/abs/2306.15794.2306.15794.

[10] Li, F., Lu, W. & Bai, Y. Hyenacircle: a hyenadna-based pretrained large language model for long eccdna prediction. Frontiers in Genetics 16, 1641162 (2025).

[11] Schiff, Y. et al. Caduceus: Bi-directional equivariant long-range dna sequence modeling. In International Conference on Machine Learning, 43632–43648 (PMLR, 2024).

[12] Liu, Z., Li, J. & Zhang, Z. eccDNAMamba: A pre-trained model for ultra-long eccDNA sequence analysis. In ICML 2025 Generative AI and Biology (GenBio) Workshop (2025). URL https://openreview.net/forum?id=56xKN7KJjy.

[13] Sennrich, R., Haddow, B. & Birch, A. Neural machine translation of rare words with subword units. arXiv preprint arXiv:1508.07909 (2015).

[14] Gage, P. A new algorithm for data compression. The C Users Journal 12, 23–38 (1994).

[15] Dao, T. & Gu, A. Transformers are ssms: Generalized models and efficient algorithms through structured state space duality. arXiv preprint arXiv:2405.21060 (2024).

[16] Zhong, T. et al. eccdna atlas: a comprehensive resource of eccdna catalog. Briefings in Bioinformatics 24, bbad037 (2023).

[17] Osama, M., Ali, S., Binte Nawaz, A., Ullah, U. et al. “extrachromosomal circular dna (eccdna): A key driver of tumorigenesis, drug resistance, and prognosis across gastrointestinal malignancies”. Health Sciences Review 15, 100228 (2025). URL https://www.sciencedirect.com/science/article/pii/S2772632025000200.

[18] Fang, M. et al. eccdna-pipe: an integrated pipeline for identification, analysis and visualization of extrachromosomal circular dna from high-throughput sequencing data. Briefings in Bioinformatics 25, bbae034 (2024). URL 10.1093/bib/bbae034. https://academic.oup.com/bib/article-pdf/25/2/bbae034/56663657/bbae034.pdf.

[19] Chang, K.-L. et al. Short human eccdnas are predictable from sequences. Briefings in Bioinformatics 24, bbad147 (2023). URL 10.1093/bib/bbad147. https://academic.oup.com/bib/article-pdf/24/3/bbad147/50410894/bbad147.pdf.

[20] Ji, Y., Zhou, Z., Liu, H. & Davuluri, R. V. Dnabert: pre-trained bidirectional encoder representations from transformers model for dna-language in genome. Bioinformatics 37, 2112–2120 (2021).

[21] Zhao, Y., Yu, L., Zhang, S., Su, X. & Zhou, X. Extrachromosomal circular dna: Current status and future prospects. Elife 11, e81412 (2022).

[22] Joshi, M. et al. Spanbert: Improving pre-training by representing and predicting spans. Transactions of the association for computational linguistics 8, 64–77 (2020).

[23] Steinegger, M. & Söding, J. Clustering huge protein sequence sets in linear time. Nature communications 9, 2542 (2018).

[24] The ENCODE Project Consortium, Moore, J. E., Purcaro, M. J. et al. Expanded encyclopaedias of dna elements in the human and mouse genomes. Nature 583, 699–710 (2020).

[25] Smit, A. F. A., Hubley, R. & Green, P. Repeatmasker open-4.0. http://www.repeatmasker.org (2013–2015).

[26] Kraft, K. et al. Enhancer activation from transposable elements in extrachromosomal dna. Nature Cell Biology (2025).

[27] Grant, C. E. & Bailey, T. L. Xstreme: Comprehensive motif analysis of biological sequence datasets. bioRxiv (2021).

[28] Gupta, S., Stamatoyannopoulos, J. A., Bailey, T. L. & Noble, W. S. Quantifying similarity between motifs. Genome Biology 8, R24 (2007).

[29] Weirauch, M. T. et al. Determination and inference of eukaryotic transcription factor sequence specificity. Cell 158, 1431–1443 (2014).

[30] Hu, X., Li, J., Fu, M., Zhao, X. & Wang, W. The jak/stat signaling pathway: from bench to clinic. Signal Transduction and Targeted Therapy 6, 402 (2021).

[31] Seachrist, D. D., Anstine, L. J. & Keri, R. A. Foxa1: A pioneer of nuclear receptor action in breast cancer. Cancers 13, 5205 (2021).

[32] Mullen, J., Kato, S., Sicklick, J. K. & Kurzrock, R. Targeting arid1a mutations in cancer. Cancer Treatment Reviews 100, 102287 (2021).

